# Community-driven shark monitoring for informed decision making: A case study from Fiji

**DOI:** 10.1101/2022.08.31.505463

**Authors:** CA Ward-Paige, H Sykes, GJ Osgood, J Brunnschweiler

## Abstract

**Context:** Globally, more than 121 million people enjoy nature-based marine tourism, making it one of the largest marine industries. Ocean degradation threatens this industry and management has not kept pace to ensure long-term sustainability. In response, some individuals within the industry are taking it upon themselves to monitor the ocean and provide the data needed to assist management decisions. Fiji is one such place.

**Aims:** Between 2012 and 2016, 39 Fijian dive operators, in collaboration with eOceans, conducted the Great Fiji Shark Count (GFSC) to document sharks on their dives.

**Methods:** Using 146,304 shark observations from 30,668 dives, we document spatial and temporal patterns of eleven shark species at 592 sites.

**Key results:** Sharks were observed on 13,846 dives (45% of recorded dives) at 441 (74%) sites. Generally, our results matched those from other, more limited surveys, including from BRUVs. We found high variability in shark presence, species richness, and relative abundance through space and time. One trend was surprising: the most common species, Whitetip Reef Shark, decreased over the study period at eastern sites and increased at western sites — the cause is currently unknown.

**Conclusions:** Our results can guide management and conservation needs, future scientific questions, and provide a baseline for future assessments.

**Implications:** This study demonstrates the value of longitudinal observation data that includes absences for describing marine fauna, and confirms the capacity of stakeholders to document the ocean. It also points the direction for broadscale participatory science methodologies to track the ocean.

## Introduction

The ocean is in a state of rapid change due to climate destabilization, acidification, and biodiversity loss (e.g., Oliver *et al*. 2018; Talukder *et al*. 2022), the impacts of which are compounded by the addition of pollutants into an ocean that has suffered decades of large-scale misuse (Halpern *et al*. 2008, 2015). In response, some international programs have been initiated to better understand and protect the ocean and the people that depend on them (e.g., The Census of Marine Life, the International Plan of Action for Conservation and Management of Sharks (IPOA-Sharks), the Convention of Biological Diversity (CBD), Aichi Targets, and the United Nations Sustainable Development Goals).

The science needed to implement, evaluate, or improve these programs pose significant challenges. Importantly, they tend to be costly and are typically ‘by invite only’. For example, the Census of Marine Life, a 10-year international effort, cost $650 million and involved only 2,700 scientists. Other stake- and rightsholders are typically not involved in these programs. However, this is slowly changing as the importance of participatory and co-generated science has come to light (Ward-Paige, Mora, *et al*. 2010; Ward-Paige *et al*. 2013; Hind-Ozan *et al*. 2017). Many scientists and managers now advocate for the inclusion of stake- and rightsholders in marine science and conservation (Cigliano and Ballard 2017), including in meeting SDGs (Fritz *et al*. 2019).

Under the broad umbrella of participatory science (e.g., citizen science, crowdsourcing, co-generated science), marine tourism can be an important partner. Individuals in the marine tourism sector regularly visit various coastal and marine ecosystems and encounter many species or threats. A few participatory science programs have already filled important data gaps on species distributions and threats, climate change induced range extensions, and fisheries, and been found to provide opportunities to promote trust, education, outreach, awareness, and best practices for ecotourism (Ward-Paige *et al*. 2014; Lawrence *et al*. 2016; Hind-Ozan *et al*. 2017). Additionally, marine tourism operators, guides, and tourists are proving to be highly motivated to document their ocean (e.g., species and anthropogenic threats) and leverage the economic value of their industry towards improved science, management, and conservation for the ocean and their livelihoods (Brunnschweiler *et al*. 2014; Ward-Paige *et al*. 2018).

Given that nature-based coastal and marine tourism has significant social and economic value as part of a growing blue economy (Cisneros-Montemayor *et al*. 2021), there is economic rationale to manage the ocean for reasons beyond commercial fishing. Globally, more than 121 million people take part in nature-based ocean activities, such as scuba diving, snorkeling, recreational fishing, and wildlife watching (Spalding *et al*. 2016). This sector generates more than $400 billion dollars per year (Spalding *et al*. 2017), rivaling commercial fisheries, aquaculture, or oil and gas in some areas (NOAA 2019). The value varies by region and market segmentation (Beaver and Keily 2015), and ecosystem (e.g., coral reefs at $37.8 billion (Spalding *et al*. 2016), but combines into one of the largest and most valued industries (Spalding *et al*. 2016, 2017). Specific species also drive these industries. For example, shark, ray, and turtle tourism attracts millions of people (e.g., scuba divers), generating direct revenues for local operators and businesses, and contributing to economies on regional and nationwide scales (Troeng and Drews 2004; O’Malley *et al*. 2013; Huveneers *et al*. 2017).

In coastal and marine environments, participatory science has been promoted as an important part of governance and management (Ward-Paige *et al*. 2014) and sustainable tourism (Lawrence *et al*. 2016). Many scientific insights have been gained because of these collaborations. For example, public contributed data have been used to document insight into ephemeral events that are hard to predict (Lester *et al*. 2022), species range extensions due to climate change (Last *et al*. 2011), exotic species invasions (Côté *et al*. 2013), large-scale absence of reef sharks in proximity to humans (Ward-Paige, Mora, *et al*. 2010), and the spatial extent of marine garbage (Jambeck and Johnsen 2015; van der Velde *et al*. 2017).

In Fiji, a substantial proportion of visitors specifically visit the country to dive with sharks, which has been estimated to input over USD 42 million annually into Fiji’s economy (Vianna *et al*. 2011; Mangubhai *et al*. 2019). Despite the considerable socioeconomic value of the diving industry, a paucity of information on the diversity, occurrence, and relative abundance of sharks remains and the community has voiced particular concern for sharks in the region (HS, JB personal communications). To address these data gaps and growing conservation interest, the Great Fiji Shark Count (GFSC) was launched in 2012 in collaboration with eOceans (previously eShark) and together with the dive tourism industry to establish the country’s first nation-wide snapshot of shark abundance and diversity. This initiative was timely with other initiatives in Fiji that have been started or pursued in earnest in the past decade, such as support for locally managed marine protected areas (Govan *et al*. 2008; Jupiter *et al*. 2014), mitigating climate change threats (Wendt *et al*. 2018), voluntary commitments to expand marine protected areas (WCS, 2017), and the adoption of a comprehensive Shark and Ray Conservation Regulation (Ministry of Fisheries and Department of Environment, Fiji). Here, we present the results from the GFSC that provide baselines of Fiji’s shark populations, including spatial and temporal trends in composition, species richness, abundance, and frequency of occurrence. Information that is critical for making science-based decisions in this Pacific Small Island Developing State.

## Methods

### Data collection

From 2012 to 2016, 39 dive operators across Fiji conducted the first nationwide longitudinal visual census of dive sites for sharks as part of the Great Fiji Shark Count (GFSC) in collaboration with eOceans (eOceans.co). Each April and November, guests (scuba divers and snorkelers) and staff from participating dive operators recorded the details of every dive at 592 sites in 25 areas across Fiji (Fig. 1) into community logbooks. Before each dive, dive guides instructed guests about the marine region, the objectives of the GFSC, and presented a field guide to correctly identify the sharks they can potentially encounter in Fiji. After the dive, each participating guest logged various attributes of the dive with their observations. For each dive, including all replicates (i.e., multiple peoples’ observations on the same site at the same time), details included: date, time in and out of the water, operator name, site name, maximum dive depth, yes or no to spearfishing, and yes or no to wildlife direct feeding or intentional attracting (Meyer *et al*. 2021). Participants recorded the presence or absence of sharks, and for each species, the number of individuals and if they observed mating or numerous juvenile sharks as an indication of potential nurseries (Heupel *et al*. 2007). To gather the most complete dataset on sharks that included zeros, and to encourage ongoing participation while gathering more information about other species, participants also recorded observations of rays, turtles, seahorses, and cetaceans (not shown here).

**Figure 1.**
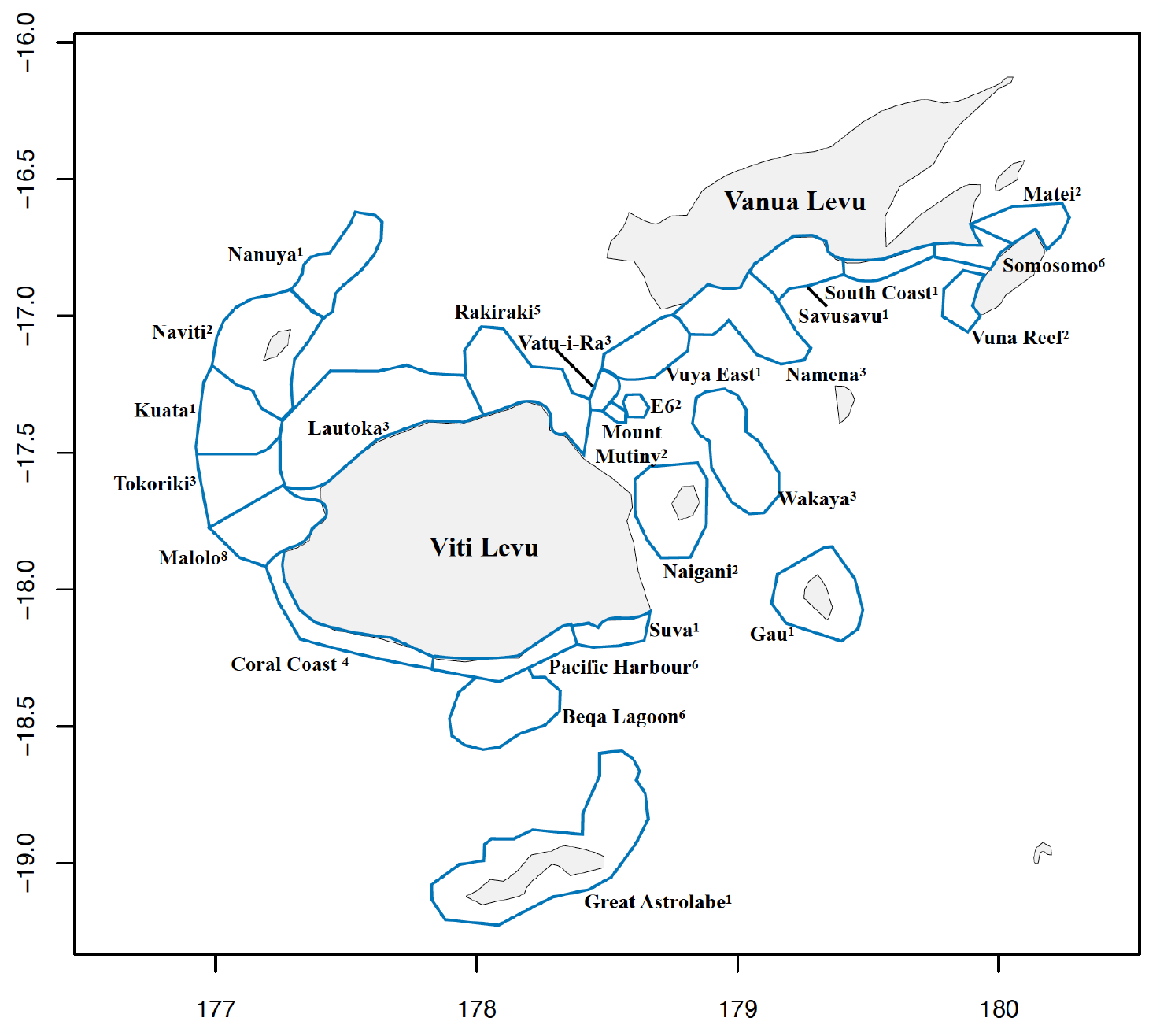
Study region with survey areas sampled by dive operators. Superscripts show the number of operators sampling each area (note: some sampled multiple areas).

### Data treatment

To protect sites, species, and local communities (e.g., from illegal fishing) we did not gather exact site coordinates and instead assigned each dive site to one of 25 areas as defined by HS’s local knowledge of how local communities use and define the areas (Fig 1). After the first year of the survey, we created a master site list for each area to accommodate differences in site nomenclature between operators. A total of 207 records were excluded because they could not be matched with a specific site.

To establish a clear picture of effort we examined and mapped the spatial and temporal patterns of survey effort (number of dives) for each site and area and controlled for variations in all analyses as described in the models section below. Additionally, we separated feeding from non-feeding sites. Ten sites in 8 areas were exclusive feeding sites. Shark feeding also occurred at 5 other sites in 2 areas but on less than 40% of the survey dives and for only short durations (e.g., one week in one year), hence we classified them as non-feeding sites. Depth was missing from 4,315 records, so we filled missing values using the average maximum depth reported at each site as a proxy. Since 31 sites (154 observations from 30 non-feeding sites and 5 observations from one feeding site) did not have any depth information, we excluded them from the statistical modeling but included them in the exploratory analyses. Additionally, five non-feeding dives lacked date information and were excluded from the statistical modeling. A total of 2,830 records (9%) at 121 sites were snorkeling surveys and were included in all analyses with other dives as our exploratory analyses did not suggest separate treatment was warranted.

### Metrics and composition analysis

Our analyses focused on the spatial and temporal distribution of sharks at feeding and non-feeding sites using three metrics: i) species richness (number of species observed per dive), ii) relative abundance (counted or estimated number of sharks per species per dive; abundance hereon), and iii) frequency of occurrence (proportion of dives a species was observed). We also investigated the composition (i.e., a metric of species evenness that includes richness and abundance) of sharks in each area, where the maximum number counted in that area at any time was used as species’ abundance, and then divided that by the total abundance across all species.

First, we summarized the spatial variation of species composition and the three metrics by area and feeding activity (yes/no). Then, to account for variations in survey effort across space and time we used generalized linear mixed models (GLMMs). We did this for the three metrics and six most common species (i.e., response variables), as determined by count and frequency of occurrence at feeding and non-feeding sites, all sites, and all areas.

For the model predictors, we set out to study both temporal and spatial trends, so we included year and month as fixed effects and dive site nested within area as random effects. For dives at non-feeding sites, random slopes for the year effect were included for both dive site and area when supported by the variability of the data (i.e., the standard deviation of the random effect was larger than 0.01) and minimized AIC. For dives at feeding sites, since regular feeding activity occurred primarily at two sites across all years (Bistro and Shark Reef Marine Reserve) in the same area (Pacific Harbour on the southern coast of Viti Levu; Fig. 1), we focused on these two sites and removed the random effects associated with dive site and area while adding a dive site by year interaction in the fixed effects when the effect was significant.

To control for variation in bottom depth and the presence of feeding activity, including at non-feeding sites, we included depth and feeding (yes/no) as fixed effects. Since there were often multiple observers on the same dive, we controlled for those correlations by including dive replicate (i.e., same dive, operator, and day) as random intercepts. For all models, Wald’s Z-tests were used to assess significance. Models were fit with restricted maximum likelihood.

Since Scalloped Hammerhead Sharks were only seen on non-feeding dives at 19 sites, their models only included those sites (a total of 3,935 dives) and did not include feeding as a fixed effect. Bull Sharks were primarily seen on feeding dives and therefore only modeled for those dives. Feeding was not included in models of Tawny Nurse Sharks at feeding sites or for Blacktip Reef Sharks at non-feeding sites due to low sample sizes.

For model distributions, the choice of Poisson over Negative Binomial and zero-inflation over none was done using AIC (delta AIC less than 3). For the three metrics, mean species richness per dive was modeled with a Poisson GLMM, mean abundance with a zero-inflated Negative Binomial GLMM, and frequency of occurrence with a Binomial GLMM. For the six species with sufficient presence data to fit a model (i.e., greater than 1000 total count and over 400 occurrences in the dataset), abundance was modeled with the following distributions: Poisson for Whitetip Reef, Blacktip Reef, and Tawny Nurse Sharks; and Negative Binomial for Grey Reef, Scalloped Hammerhead, and Bull Sharks. All models included zero inflation except for Whitetip Reef and Blacktip Reef Sharks, which were not zero-inflated.

A note for added clarity regarding replicates. Our analyses assumed all observations are replicates whether on the same dive, day, or week. The observations were not summed across replicates but the replicates rather inform the averages for occurrence and abundance.

## Results

The GFSC — a collaborative endeavor of dive operators, dive masters and their guests, with scientists — achieved extensive survey effort over space and time (Fig. 2). In total, 27,838 dive and 2,830 snorkeling (dives hereon) logs were analyzed from 592 sites (577 = strictly non-feeding, 6 = exclusively feeding, 9 = both) in 25 areas in April and November over five years. Eleven shark species from 4 families (Carcharhinidae, Sphyrnidae, Ginglymostomatidae and Stegostomatidae) and 2 orders (Carcharhiniformes and Orectolobiformes) were reported, all of which have a Red List status of Near Threatened or worse, with populations either decreasing or unknown trends (Table 1). Mating was reported on 32 dives (<1%) from 7 sites (1%), juveniles on 1,355 dives (4%) from 23 sites (4%; Table S1), and spearfishing on 16 dives (<1%) at 2 sites.

**Figure 2.**
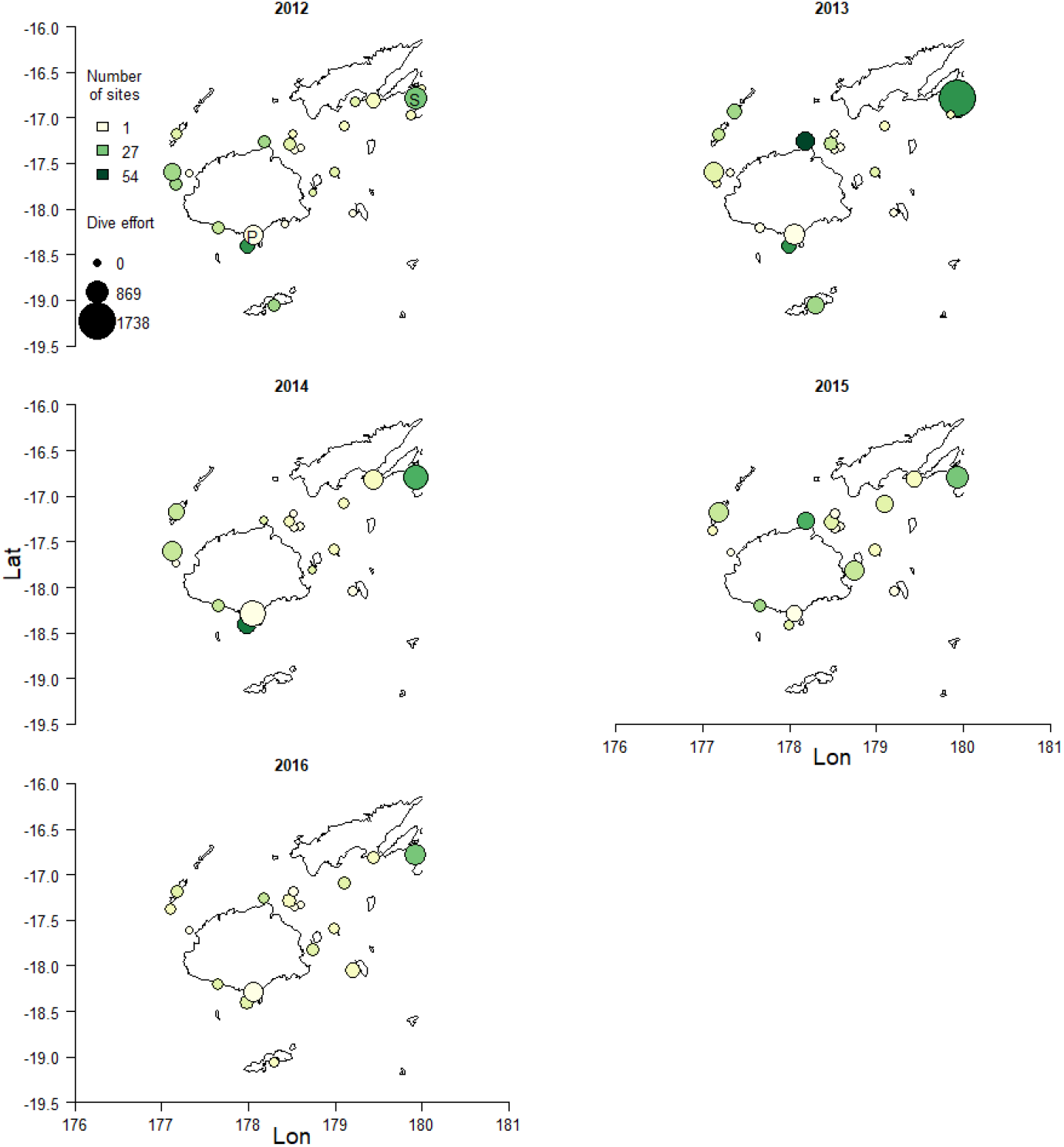
Survey effort through space and time across all dives. Point size scales to number of dives and point color to number of sites visited in each area. P = represents Pacific Harbour and S = Somosomo Straits (in 2012 panel).

**Table 1.** Summary of shark species encountered in the Great Fiji Shark Count. Shark species encountered with respective mean and maximum abundance per dive, mean school (group) size (mean count excluding zeros), and frequency of occurrence during both non-feeding and feeding dives. Red List Status according to the International Union for Conservation of Nature (IUCN) is NT = Near Threatened; VU = Vulnerable; EN = Endangered; CR = Critically Endangered.

| Common name | Latin name | IUCN Red List Status | Mean abundance |  | Maximum abundance |  | Mean school (group) size |  | Frequency of occurrence |  |
| --- | --- | --- | --- | --- | --- | --- | --- | --- | --- | --- |
|  |  |  | Non-feeding | Feeding | Non-feeding | Feeding | Non-feeding | Feeding | Non-feeding | Feeding |
| Bull Shark | *Carcharhinus leucas* | NT (Unknown) | 0.003 | 14.2 | 20 | 100 | 3.6 | 16 | 0.00082 | 0.87 |
| Whitetip Reef Shark | *Triaenodon obesus* | NT (Unknown) | 0.63 | 4.55 | 26 | 60 | 2 | 8.5 | 0.32 | 0.53 |
| Grey Reef Shark | *Carcharhinus amblyrhynchos* | NT (Unknown) | 0.55 | 3.38 | 88 | 41 | 5.6 | 6.1 | 0.099 | 0.55 |
| Blacktip Reef Shark | *Carcharhinus melanopterus* | NT (Unknown) | 0.028 | 3.30 | 9 | 85 | 1.4 | 7.9 | 0.019 | 0.41 |
| Tawny Nurse Shark | *Nebrius ferrugineus* | VU (Decreasing) | 0.0048 | 0.86 | 3 | 40 | 1.1 | 4 | 0.0043 | 0.21 |
| Scalloped Hammerhead Shark | *Sphyrna lewini* | CR (Decreasing) | 0.40 | 0 | 100 | 0 | 23 | NA | 0.017 | 0 |
| Sicklefin Lemon Shark | *Negaprion acutidens* | VU (Decreasing) | 0.00019 | 0.23 | 2 | 10 | 1.2 | 2.8 | 0.00015 | 0.083 |
| Silvertip Shark | *Carcharhinus albimarginatus* | VU (Decreasing) | 0.0044 | 0.13 | 24 | 12 | 1.6 | 2.4 | 0.0028 | 0.056 |
| Zebra Shark | *Stegostoma fasciatum* | EN (Decreasing) | 0.0097 | 0.0018 | 5 | 2 | 1.6 | 1.2 | 0.006 | 0.0016 |
| Tiger Shark | *Galeocerdo cuvier* | NT (Decreasing) | 0.00071 | 0.011 | 2 | 5 | 1.1 | 1.1 | 0.00067 | 0.0097 |
| Great Hammerhead Shark | *Sphyrna mokarran* | CR (Decreasing) | 0.00071 | 0 | 1 | 0 | 1 | NA | 0.00071 | 0 |

### Survey effort

Effort was consistent across years (5,139 to 7,018 dives) and months (15,283 in April to 15,385 in November), but varied by area (12 to 5,359 dives, >50% of areas had >1000 dives; Fig 2). Areas also varied in number of operators (1 to 8) and sites (1 to 82), where Somosomo had the highest total effort with 5,359 dives (all non-feeding) followed by Pacific Harbour with 3,918 dives, 3,500 of which were feeding dives (Fig 2, Table S1). Feeding occurred on 3,816 dives (12%) at 15 sites (2%) in 10 areas (40%), but feeding only regularly occurred at 10 sites in 8 areas. The majority of feeding dives occurred at 2 sites in Pacific Harbour, Shark Reef Marine Reserve (81.7%) and Bistro (10.4%) (Table S1, S2).

### Spatial patterns

Sharks were reported in all areas with mean species richness, abundance, and frequency of occurrence varying by area (Fig. 3, 4). In total, 146,304 recordings of sharks from 13,846 dives (45% of recorded dives) at 441 (74%) sites were reported (Table S1). The number of species observed generally increased with survey effort to a maximum of 9 species reported from Pacific Harbour (Fig. S1). Overall, species richness and abundance was higher at feeding sites compared to non-feeding sites (Fig. 3 a,c, Fig. S2),

**Figure 3.**
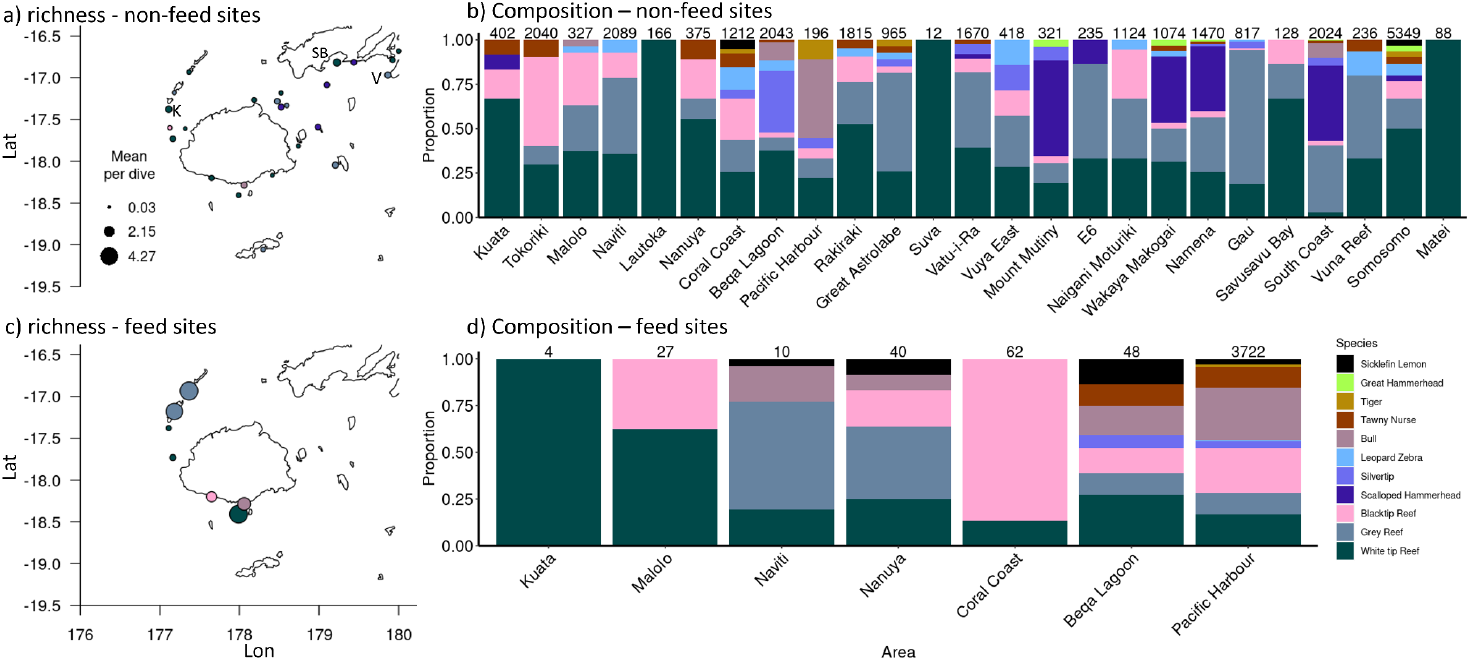
Shark species composition by area at non-feeding (top) and feeding (bottom) sites. Shown is mean species richness by area (a, c) and the proportion of all shark observations made by each species at each area, ordered from west to east (b, d). Point color on maps reflects the most commonly observed species, and numbers above bars are number of dives. SB = Savusavu Bay, V = Vuna Reef, K = Kuata (in a) panel).

**Figure 4.**
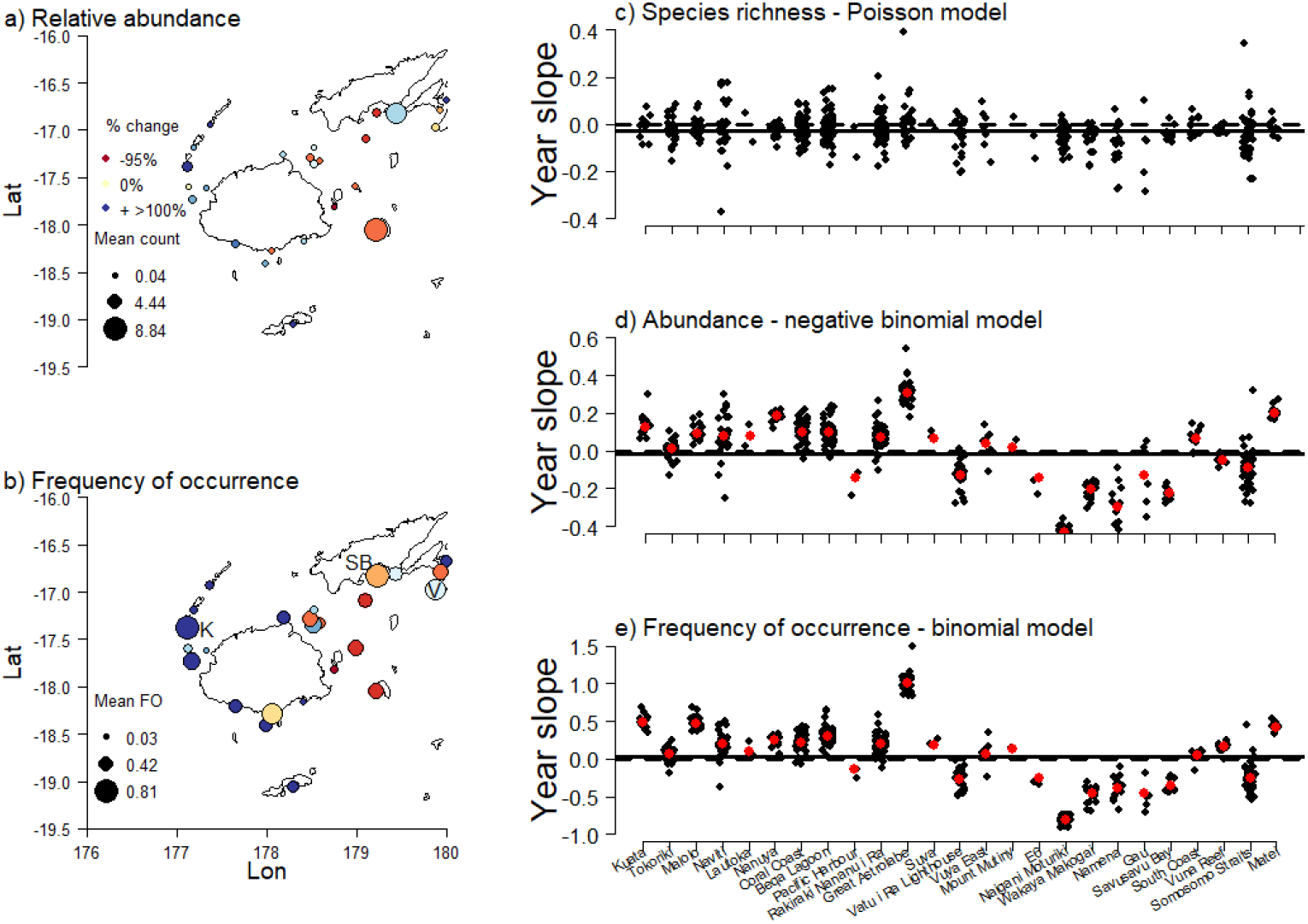
Change in species composition by area and year, including mean species richness, abundance, and frequency of occurrence at non-feeding dives. Maps of (a) abundance and (b) frequency of occurrence by area where point color indicates percent change predicted from GLMMs between 2012 and 2016 and point size indicates average count or frequency of occurrence by area averaged over all years and sites. Species richness variation did not support random slopes by area. The slope of the year effect, accounting for random effects, for each site within an area (black) and for the area when present (red) for the species richness Poisson model (c), the abundance negative binomial model (d), and the frequency of occurrence binomial model (e). The solid line represents the overall slope of the year effect marginalized across area. The dotted line represents zero. Sites are ordered left to right by west to east. SB = Savusavu Bay, V = Vuna Reef, K = Kuata (in b) panel).

#### Non-feeding sites

Mean species richness varied little by area (Fig. 3a) and over 82% of dive sites (88% of those dived over 100 times) had less than one species observed per dive on average. Savusavu, followed by Kuata, Gau, and Vuna Reef consistently reported the most shark species at non-feeding sites, with mean richness around one species (Fig. 3a, Table S2).

Mean abundance was generally low but varied by area from <0.1 sharks of any species per dive in Lautoka to 9 sharks in Gau (Fig. 4a, Table S1, S2). Most non-feeding areas (92%) and sites (96%) reported less than three sharks of any species per dive on average, and 48% of areas and 74% of sites reported less than one (Fig. 4a; Table S1, S2). On average, school (group) size (mean abundance excluding zeros) was 4 sharks of any species per dive (range 1 to 125). The majority (87%) of sightings were individuals or small groups with <5 individuals; the largest schools (>50 sharks) were of Scalloped Hammerhead and Grey Reef Sharks, primarily at Dream House and Nigali Passage, respectively (Table S2).

Frequency of occurrence varied greatly. Sharks were most frequently (>70%) observed in the Savusavu, Kuata, and Vuna Reef, and least (<10%) in the Lautoka and Suva areas (Fig. 4b, Table S1). Frequency of occurrence was greater than 90% at 79 non-feeding sites (Table S2).

Species composition varied by area. The three most abundant and frequently observed species at non-feeding sites were Whitetip Reef, Grey Reef and Blacktip Reef Sharks (Fig. 3b). Whitetip Reef and Grey Reef Sharks made up the majority of individual sharks observed at non-feeding sites: on 32% and 10% of dives, with mean group sizes of 2 and 5.6 sharks, respectively, while all other species were seen on less than 2% of dives (Table 1). The largest schools (groups) were formed by Scalloped Hammerhead and Grey Reef Sharks with mean school sizes of 23 and 5.6 sharks, and maximum number of animals observed together of 100 and 88 individuals, respectively (Table 1, S2).

#### Feeding sites

Shark encounters at feeding sites were frequent and abundant with 26 sharks being reported on average per dive compared to less than two sharks from non-feeding sites (Fig. S2). Most sharks were observed in the Pacific Harbour and Coral Coast areas with mean shark counts ranging from 21 to 25 sharks per dive, respectively. Across all species, mean school (group) size (i.e. excluding zeros) was 28 individuals on feeding dives, with the largest groups (30.19 ± 0.08) observed at the Shark Reef Marine Reserve. For sites surveyed over 100 times, the highest mean species richness and frequency of occurrence were recorded at Bistro and Shark Reef Marine Reserve in the Pacific Harbour area with shark encounters on >95% of dives and mean species richness of 2.5 and 4.5 species per dive, respectively (Table S2).

In contrast to non-feeding sites, Bull Sharks were the most commonly observed and abundant species on feeding sites (Table 1), and similar to non-feeding sites, Grey Reef and Whitetip Reef Sharks were also common (Table 1, Fig. 3d). Bull Sharks occurred on 87% of all feeding dives with group sizes averaging 16 sharks, followed by Grey Reef Sharks on 55% of dives with an average group size of 3.4. Whitetip Reef Sharks occurred on 53% of dives with an average group size of just under 9. Feeding attracted a smaller abundance (mean abundance <8) of Blacktip Reef Sharks per dive, but they were still present on 41% of dives and in most areas where feeding occurred (Fig. 3d). The largest schools were formed by Bull Sharks, with groups of up to 100 individuals; Grey Reef, Whitetip Reef, Blacktip Reef, and Tawny Nurse Sharks also occasionally formed large schools at feeding sites (Table 1).

Feeding sites in the Pacific Harbour area had the highest encounter rates for Grey Reef, Blacktip Reef, Bull, and Tawny Nurse Sharks, as well as Silvertip and Sicklefin Lemon Sharks (Table S2). Bull Sharks were commonly and abundantly observed at the feeding sites in Pacific Harbour, Beqa Lagoon, and Naviti areas (Fig. 3d). In general, all species except Zebra, Scalloped, and Great Hammerhead Sharks were more frequent and abundant on average at feeding compared to non-feeding sites (Table 1, Fig. 3c).

### Shark temporal patterns

#### Non-feeding sites

Across Fiji, there was no strong interannual trend in mean species richness, abundance, or frequency of occurrence of sharks at non-feeding sites, but there was variation at the area and site levels (Table 2, Fig. 4c,d,e, Fig. S3–S5). Each metric was higher — statistically significant for species richness and abundance — in April compared to November (Table 2). Increases in species richness occurred at 27% of sites (Fig. 4c), and most dive sites and areas (60–67%) also showed an increase in shark abundance (Fig. 4d) and frequency of occurrence (Fig. 4e) between 2012 and 2016, particularly in the west of Fiji (Fig. 4a,b). On average, sites in Naigani, Wakaya, and Namena saw the largest declines across all metrics, up to 95% for frequency of occurrence (Fig. 4a,b, Table S2). Generally, areas and sites with more negative interannual trends had a higher starting abundance, as the random intercept and slope had strong (<-0.50) negative correlations in all models.

**Table 2.** Factors influencing species richness, abundance and frequency of occurrence. Coefficients (and p-values based on Wald’s Z-test) for each effect from each model as well as the variance of the area and dive site random slopes and intercepts when fit. Feeding compares feeding and non-feeding dives at the same site. *Feeding models only used data from feeding sites in the Pacific Harbour area, therefore the dive site intercept and effect refers to Bistro compared to Shark Reef Marine Reserve (to which the main effects refer).

| Model | Year | Month:<br>November | Depth | Feeding<br>dives | Area:<br>intercept | Area: year<br>effect | Dive site:<br>intercept* | Dive site: year<br>effect* |
| --- | --- | --- | --- | --- | --- | --- | --- | --- |
| **Non-feeding sites** |  |  |  |  |  |  |  |  |
| Richness | -0.027<br>(0.30) | **-0.12**<br>**(0.0050)** | **0.064**<br>**(0.0095)** | -0.11<br>(0.72) | 0.46 | – | 1.08 | 0.041 |
| Abundance | -0.013<br>(0.85) | **-0.12 (0.040)** | **0.20**<br>**(**<0.001**)** | -0.062<br>(0.88) | 1.41 | 0.064 | 1.85 | 0.051 |
| Frequency of occurrence | 0.037<br>(0.86) | -0.22 (0.15) | 0.11 (0.21) | 0.49<br>(0.75) | 9.89 | 0.65 | 12.03 | 0.30 |
| Whitetip Reef Shark | -0.017<br>(0.82) | **-0.15 (0.013)** | 0.049 (0.17) | -0.078<br>(0.86) | 1.45 | 0.078 | 1.93 | 0.051 |
| Grey Reef Shark | 0.013<br>(0.87) | 0.15 (0.27) | **0.65**<br>**(**<0.001**)** | – | 3.46 | – | 7.39 | 0.14 |
| Blacktip Reef Shark | -0.27 (0.26) | 0.12 (0.64) | -0.26 (0.14) | – | 6.22 | 0.73 | 9.00 | 0.81 |
| Scalloped Hammerhead Shark | 0.12 (0.54) | -0.025 (0.96) | **2.21**<br>**(**<0.001**)** | – | – | – | 5.83 | – |
| Tawny Nurse Shark | -0.053<br>(0.83) | 0.42 (0.52) | -0.066 (0.88) | 2.68<br>(0.35) | – | – | 12.24 | – |
| **Feeding sites** |  |  |  |  |  |  |  |  |
| Richness | 0.018<br>(0.19) | **-0.093**<br>**(0.0084)** | **0.48**<br>**(**<0.001**)** | **0.52**<br>**(**<0.001**)** | – | – | **0.32 (0.0030)** | **0.19 (**<0.001**)** |
| Abundance | 0.02 (0.35) | **-0.73**<br>**(**<0.001**)** | **0.29**<br>**(**<0.001**)** | **2.22**<br>**(**<0.001**)** | – | – | **-0.66**<br>**(**<0.001**)** | **0.26 (**<0.001**)** |
| Frequency of occurrence | **0.59**<br>**(**<0.001**)** | 0.49 (0.15) | 0.052 (0.81) | **2.92**<br>**(**<0.001**)** | – | – | 0.52 (0.32) | – |
| Whitetip Reef Shark | **-0.36**<br>**(**<0.001**)** | **-0.97**<br>**(**<0.001**)** | **2.45**<br>**(**<0.001**)** | **-1.93**<br>**(**<0.001**)** | – | – | **-2.26**<br>**(**<0.001**)** | **1.26 (**<0.001**)** |
| Grey Reef Shark | 0.070<br>(0.18) | 0.22 (0.10) | **1.74**<br>**(**<0.001**)** | **0.79**<br>**(0.033)** | – | – | **-1.37**<br>**(0.0014)\*** | **0.76 (**<0.001**)\*** |
| Blacktip Reef Shark | -0.14<br>(0.091) | **-0.63**<br>**(0.0041)** | **2.61**<br>**(**<0.001**)** | 1.21<br>(0.10) | – | – | **-2.36**<br>**(**<0.001**)** | **0.75 (0.0041)** |
| Bull Shark | 0.049<br>(0.081) | **-0.79**<br>**(**<0.001**)** | **-0.18**<br>**(**<0.001**)** | **3.49**<br>**(**<0.001**)** | – | – | **-1.34**<br>**(**<0.001**)** | **0.23 (0.0048)** |
| Tawny Nurse Shark | 0.058<br>(0.56) | **-2.16**<br>**(**<0.001**)** | 0.011 (0.49) | – | – | – | **5.73 (**<0.001**)** | – |

At the species level, there were no statistically significant trends in abundance across Fiji (Table 2), but there was significant variation in trends by area and site for Whitetip Reef, Blacktip Reef, and Grey Reef Sharks (Fig. 5), the only species abundant enough for a random slope by site to be fit. Whitetip Reef Sharks showed >20% decreases and increases in 11 (44%) and 12 (48%) areas, respectively (Fig. 5a,d). The largest declines occurred in the east of Fiji, particularly in the Naigani (−91%), Namena (−82%), Wakaya (−63%), and Gau (−50%) areas, and the greatest increases in the Great Astrolabe (+450%), Naviti (+103%), and Beqa Lagoon (+103%) areas (Fig. 5a,d). Grey Reef Sharks were the only species that had consistent increasing abundance, but the trend was not statistically significant (Table 2, Fig. 5b,e, Fig. S6b). Blacktip Reef Sharks showed declines at many sites across Fiji, but the overall decline was not significant (Table 2, Fig. 5cf, Fig. S6d). The trends of the other species fluctuated across years, but did not vary enough by site for site-specific trends to be modeled (Fig. S6). Scalloped Hammerhead and Tawny Nurse Shark observations were least variable among sites where they were observed (Table 2).

**Figure 5.**
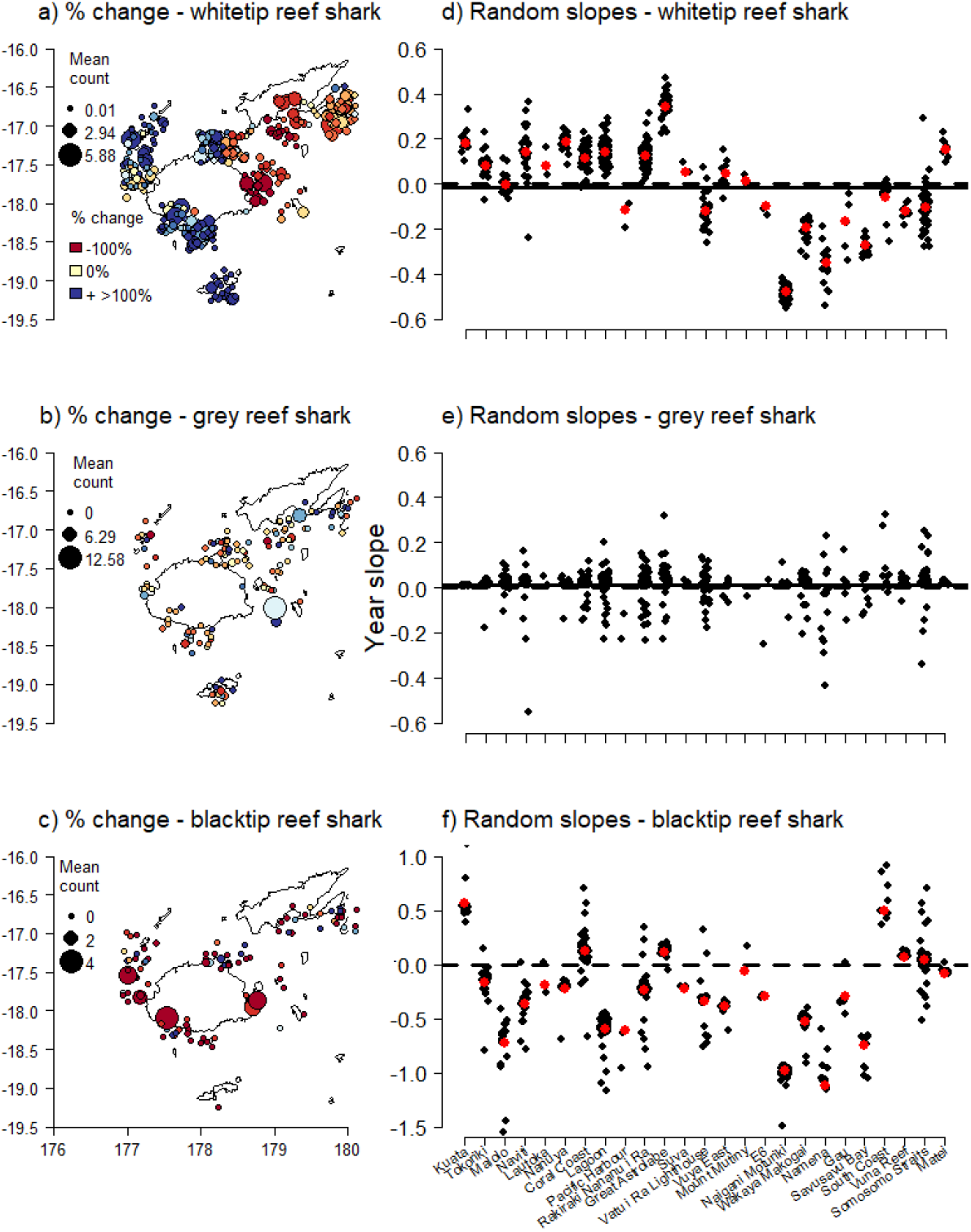
The random slope of the year effect on the abundance of (a, d) the Whitetip Reef Shark model, (b, e) the Grey Reef Shark model, and (c, f) the Blacktip Reef Shark model at non-feeding sites in each area. (a–c) Maps with site locations jittered from their area latitude/longitude, where point colour indicates percent change predicted at a site from the GLMMs between 2012 and 2016 and point size indicates average count of that species at that site. Only sites where a species was observed at least once are included. (d–f) Plots of the random slopes from the GLMMs. Red dots represent the area-specific random slope and black dots the random slopes per dive site within area. The solid line represents the overall slope of the year effect marginalized across area. The dotted line represents zero. Sites along the x-axis are ordered from west to east.

#### Feeding sites

The two most visited feeding sites, Shark Reef Marine Reserve and Bistro in the Pacific Harbour area had fluctuating but differing trends in species richness, abundance, and frequency of occurrence (Fig. 6). Species richness was higher at Bistro, and increased throughout the study period (Fig. 6a). Shark abundance for most species was higher at Shark Reef Marine Reserve (Table 2, Fig. 6a,b, Fig. S7). Frequency of occurrence increased significantly at both sites, with a ∼179% increase in predicted frequency of occurrence at each site (Table 2, Fig. 6). However, by species, frequency of occurrence increased only for Grey Reef Sharks at Bistro (Table 2, Fig. S7c). Annual trends in Bull, Whitetip Reef, and Blacktip Reef Sharks also had trends that differed by each site (Table 2); however, the trends were small compared to their fluctuations in abundance across years (Table 2, Fig S7).

**Figure 6.**
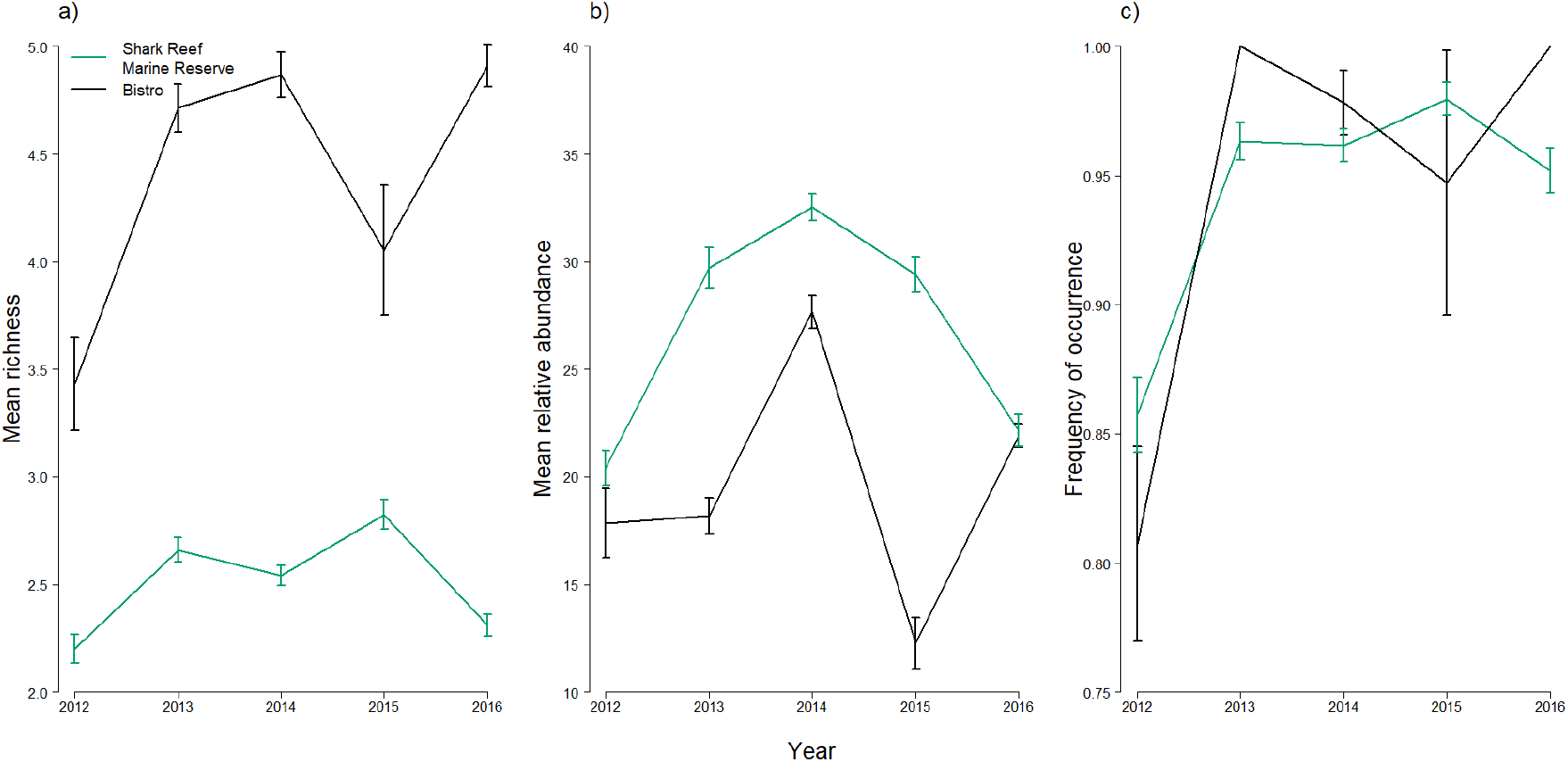
Sharks at feeding sites by year in Pacific Harbour. Annual (a) mean richness (± SE), (b) mean abundance (± SE), and (c) frequency of occurrence (± SE) at feeding sites in the Pacific Harbour area.

## Discussion

### Summary

The Great Fiji Shark Count (GFSC) — a collaboration of 39 dive operators, scientists and eOceans — conducted a nation-wide five-year census of 592 sites across Fiji by collecting observations from >30,000 dives in 25 areas. A total of 146,304 shark observations were used to describe at-sea spatial and temporal patterns of population composition, species richness, occurrence, and relative abundance. Such species’ distribution patterns are largely undescribed elsewhere in the world at this spatial and temporal scale, demonstrating the value of this type of collaborative program for revealing novel information about highly sought, mobile, and vulnerable species. All species varied in occurrence and abundance at site and area levels, demonstrating the need for high resolution longitudinal data that includes observations when no sharks were observed. Feeding elevated shark species richness, occurrence, and abundance, but the effect was site and area specific. Additionally, the long-term, ongoing contribution by dive operators to the GFSC demonstrated high-level of interest by the community to document their activities and ecosystem. Taken together, these observations suggest that some members of the dive tourism industry are motivated to track ocean issues that matter to them, which may serve the broader interests of the community, scientists, and decision makers if they find ways to collaborate to build trust, improve the speed, transparency, and accuracy of discoveries, and to make informed and accepted management and policy decisions.

### Observed patterns

By reporting every dive, regardless of what was observed (e.g., no sharks), the GFSC was able to gain novel insights on variations of sharks at the species level, which can inform and prioritize future scientific research and management.

Our study demonstrated that various species of sharks are seen throughout Fiji and that they are common and abundant enough to be detected by divers in all areas. This is positive since all species except Tiger Sharks are Threatened — two are Critically Endangered, three are Endangered, and five are Vulnerable according to the IUCN Red List of Threatened Species (https://www.iucnredlist.org/; Table 2); many have life history characteristics that make them vulnerable to exploitation (Dulvy and Forrest 2012); many have a long history of overexploitation and are still targeted (Oliver *et al*. 2015; Worm *et al*. 2013); and many live in coastal environments where they are exposed to numerous threats (Lotze *et al*. 2006; Waycott *et al*. 2009; Ward-Paige *et al*. 2015). Additionally, overexploitation and bycatch of sharks has been rampant around the world (Oliver *et al*. 2015; Worm *et al*. 2013; Wallace *et al*. 2010; Dulvy *et al*. 2014), including in Fiji (Glaus *et al*. 2015), and many species are now too rare to be detected by divers in other regions (Ward-Paige, Mora, *et al*. 2010).

#### Alignment with other studies

The spatial ecology of the majority of shark species documented in the GFSC has not been described across Fiji before. However, where there is overlap there are mostly only minor deviations in the numbers of individuals and locations. This correspondence adds some level of validation to the data collection methodologies and results as presented.

Shark species reported in the GFSC are the same as those documented by divers previously (Brunnschweiler and Earle 2006; Vianna *et al*. 2011), and only the Blacktip Shark (*Carcharhinus limbatus*; caught by fishers) (Glaus *et al*. 2018) and the Whale Shark (*Rhincodon typus*; on offshore sites in northwest Viti Levu) (Sykes *et al*. 2018) were not recorded in the GFSC. Small numbers of adult Scalloped Hammerhead Sharks have been documented in the Vatu-i-Ra Lighthouse area before (Vianna *et al*. 2011), but our study is the first to record schooling specimens of this species in Mount Mutiny, Wakaya, Namena, and South Coast of Vanua Levu. These additional and less frequented sites possibly connect the South Coast of Vanua Levu adult population to the juvenile Scalloped Hammerhead Sharks that are found in Suva Harbour (Marie *et al*. 2017), but further investigation is needed.

A 2004 scientific diver study in the Shark Reef Marine Reserve found that eight shark species use the site (Brunnschweiler and Earle 2006). The GFSC found that the same species still use the site, with the addition of Zebra Sharks, but the mean numbers appear to have changed slightly for a few species with the GFSC having higher Blacktip Reef (7.9 sharks per dive compared to ∼1) and Whitetip Reef Sharks (9.6 sharks per dive compared to ∼1), and lower numbers of Tawny Nurse (1.1 sharks per dive compared to ∼4) and Grey Reef Sharks (6.4 sharks per dive compared to ∼15) (note: monthly and annual comparisons are needed to be more explicit about change). In Namena, which contains Fiji’s second largest no-take marine reserve, a one-month study in July 2009 using baited remote underwater video systems (BRUVs) found five shark species (Goetze and Fullwood 2013); with the exception of Zebra Sharks (previously a maximum of one individual) the GFSC found the same species, with the addition of Tawny Nurse (max = 1, mean = 1), Scalloped Hammerhead (max = 30, mean = 3.8), and Great Hammerhead Sharks (max = 1, mean =1). Interestingly, for the species occurring in both studies, the maximum number of individuals seen in the previous study are very similar to the GFSC — Grey Reef at 19 and 25 (GFSC mean = 3.3), Whitetip Reef at 19 and 21 (GFSC mean = 2), Blacktip Reef at 3 and 3 (GFSC mean = 1.7), and Silvertip Sharks at 1 and 1, respectively. Further, these similarities between the GFSC diver data and BRUV data demonstrate the value of high effort dive sampling, and the lower mean values of the GFSC demonstrate the value of longitudinal data for detecting variations in populations through time. These two methods, if deployed together, may greatly increase what is known about shark populations and their conservation needs.

#### Novel findings

The most surprising result was that, over this relatively short study period, abundance of Whitetip Reef Sharks significantly increased at nearly all western sites and significantly decreased at nearly all eastern sites (Fig. 5 a,d). Rapid decreases of shark abundance at a site are possible — habitat destruction and fishing, for example, can quickly remove sharks from an area. Rapid increases at a site, on the other hand, are more challenging to explain because the sharks would either need to be born (need all elements of recruitment) or move in from elsewhere (migration). An increase in abundance can happen in a specific area when, for example, a new feed site or artificial reef is established. However, in such a case, Whitetip Reef Sharks would be expected to move in from nearby sites as studies have shown that this species has relatively small home ranges and typically moves tens of kilometers (Speed *et al*. 2011; Barnett *et al*. 2012) rather than the hundreds of kilometers it would take it to move from the eastern sites to the western sites. Another explanation might be that while the models controlled for year, month, site, area, depth, effort, and feeding, it is possible that another unknown and undocumented change to observations occurred (e.g., visibility at all the eastern sites decreased while visibility across all western sites increased). We are, however, unaware of any systematic changes affecting all sites. As we have no satisfactory explanation for this broad-scale pattern at this stage, we suggest that an in-depth co-interpretation study with the community, such as through a local knowledge survey, might bring forth suggested hypotheses for future testing.

Seasonal and annual variations at the site and area level have not been previously documented at this spatial and temporal scale and demonstrates the importance of longitudinal sampling for the study of mobile marine fauna. Our results may have important implications for scientific investigations of mobile marine megafauna more generally, especially for observational studies that have limited spatial and temporal scope. By regularly sampling sites over five years, the GFSC showed clear variations in species occurrence and abundance. Many scientific studies, especially in remote areas, are resource and time limited — covering a relatively short time period (typically only a few days or weeks) at a small number of sites, often without replicates. For mobile species, even those with relatively small home ranges like reef sharks, these small and short-term censuses may misrepresent populations (e.g., missing species). Therefore, expanding such studies through participatory monitoring could help capture the sensitivity of the sampling methodologies and add perspective on these short snapshots.

Mating and schools of juvenile sharks were only occasionally observed, and were variable through space and time. This limits our results for identifying sensitive nursery areas that are among the most important areas to be considered for shark management and conservation (Kinney and Simpfendorfer 2009). However, conducting an even higher resolution censuses, such as by running the GFSC all year, including others’ observations (e.g., fishers), and combining with other survey types (e.g., BRUVs) (Vierus *et al*. 2018), may help to better identify these areas and their variation by species.

Finally, our results marginally extend what is known about the impacts of feeding or intentionally attracting sharks in Fiji (Brunnschweiler and Baensch 2011; Brunnschweiler *et al*. 2014). Without pre-feed baselines, we cannot fully understand the impact — presumably sharks already occurred in higher abundance on these sites compared to other sites but this is not documented and is beyond the scope of this study. Regardless, feeding is an important consideration in understanding and interpreting our results and the contemporary distribution of sharks. Bull sharks, for example, are one of the most sought species for shark diving tourism in Fiji (Brunnschweiler *et al*. 2014) and were reported in six areas during the GFSC.

Bull Shark occurrence and abundance was highest at two sites in Pacific Harbour, where feeding occurred regularly. Interestingly, however, the area with the second highest abundance of Bull Sharks was in South Coast of Vanua Levu (maximum of 20 and mean of 6.6 individuals), which did not report shark feeding or attracting. It remains unknown what attracts Bull Sharks to this area and more importantly if these are the same individuals that can be observed at the SRMR in the Pacific Harbour area (Brunnschweiler and Baensch 2011). Bull sharks are capable of long-range movements (Brunnschweiler *et al*. 2010; Heupel *et al*. 2015), and individuals that were visually identified at the SRMR on the southern coast of Viti Levu were also recorded 200 km away at Yakawe Reef in the northwest of Fiji (Yasawa Group) where a shark feed site was established in 2015 (Krüger 2021). Regardless, these differences further highlight the importance of tracking and controlling for feeding when describing and monitoring marine animals.

### Caveats for observed patterns

There are some general caveats to consider. Sampling was not standardized in space or time, which limits some of the potential analyses and interpretations, but varying effort was controlled for in the models. As with all visual censuses, visibility, distance to the animal, and diver experience can affect species detection and identification (Thresher and Gunn 1986; Darwall and Dulvy 1996; Ward-Paige, Mills Flemming, *et al*. 2010) and were not addressed in this study. Some species, populations, and individuals may display more or less avoidance behavior to scuba divers (MacNeil *et al*. 2008; Cubero-Pardo *et al*. 2011) including in shark feeding areas (Brunnschweiler *et al*. 2014), but human activities are a part of the ecosystem that are always changing and need to be considered. Because the exact site locations (to protect people and species) were not reported, we cannot estimate the impact of feeding on adjacent sites — we would expect site-site proximity and species mobility to be important factors determining the independence of sites (e.g., if the same individuals are observed in nearby or distant sites and areas) — but future tagging studies and distance between sites could help refine models. When calculating abundance as the mean school size of each species, we assumed that all sightings were repeated sightings and therefore underestimated relative abundance, especially for highly mobile and transient species that are unlikely to be repeatedly observed over multiple years. Snorkeling and scuba diving observations were combined as our initial investigations did not warrant their separation (Fig S8, S9); however, future studies that track more benthic and cryptic species may need to include activity type in the models to control for viewing differences. Visibility can determine what is observed or missed, but previous investigations found too much variability in visibility classification to be precisely or accurately captured without significant training and standardization (CWP personal observation) and because these tourist-driven sites are well known for high visibility, it was not gathered in the GFSC; however, future studies that take place in areas with higher variation in visibility or sampling smaller or more cryptic species should consider including visibility in the data gathering and models. Finally, relative abundance is not expected to reflect true abundance, but rather a proxy of the number of individuals observed in each area, and these values need to be carefully considered with human use patterns that may impact shark behavior (e.g., feeding). Many of these challenges can be overcome by considering population specific information on home range, residency, and site fidelity, which could help to estimate repeat versus independent sightings and refine relative abundance estimates. We do not yet have this level of detail, but tagging studies, such as those done for Bull Sharks in the Pacific Harbour area (Brunnschweiler and Barnett 2013), may help to define this further in the future.

### Considerations for future work

This first description of the spatial patterns of eleven shark species across Fiji lays the foundation for further scientific research, conservation, and management design or evaluation for sharks in Fiji. It also establishes a framework and process for collaboratively tracking different dimensions of the ocean, including other species, areas, and evolving issues (e.g., spills, climate change, habitat destruction) affecting species and ecosystems. Pursuing projects that are important to and led by the community may be the most influential factor determining long-term participation and success. Innovating solutions for scaling these types of participatory and co-generated projects with self-determination and self-declaration tools for data sharing, knowledge transfer, with experts performing timely standardization, quality control, and analysis will be key.

Future research on sharks in Fiji could use a number of approaches to build on the GFSC framework and results presented here. For example, expanding the GFCS program to gather diver observations throughout the year and incorporating observations from other activities or stake- and rightsholder groups would increase the spatial and temporal resolution of shark distributions and could be useful for tracking higher resolution trends through space and time, refining knowledge on potential nursery sites, to inform policy and priority protection areas and strategies, and possibly enable dynamic management if the data can be collected, analyzed, and distributed in a timely fashion. Combining spatial GFSC observation data with ecosystem data, such as temperature, habitat type, depth, other species, human dimensions (e.g., development, shipping, fishing) could be used in a habitat modeling exercise to understand the drivers of the spatiotemporal trends of sharks found in the GFSC, and thus help to understand the potential impacts of change (e.g., climate change, habitat loss/restoration, fishing management). Soliciting additional data and input variables, such as photographs or records of shark condition (e.g., size, scars, parasites, entanglements, hooks) as has been done elsewhere (Bradshaw *et al*. 2007; Dudgeon *et al*. 2008; Couturier *et al*. 2011; Araujo *et al*. 2017) and could be deployed in a future GFSC project to identify priorities for evolving threats and needs for conservation and management. Additionally, complementing observation data with BRUVs, photographic mark-recapture, tagging, and tracking studies could help further refine understanding on the influence of feeding, how sharks move between sites and areas, and how they move in response to the presence of humans.

The success of the GFSC suggests that it could be a framework for tracking other species, areas, and dimensions of the ocean, which would include other ocean stakeholder groups. Success of the GFSC can be measured in many ways – community interest, inclusion of different perspectives (i.e., operators, guides, tourists), the relatively high spatiotemporal resolution of data or the number of sites surveyed. This could be instrumental for understanding marine animal populations, ecosystems, and the human dimensions of these ecosystems because of the challenging logistical nature of marine field studies that largely limit the space and duration of studies – often only sampling a handful of sites within a few weeks or months. The GFSC provides context for what is possible through participatory science processes. We suggest that if traditional sampling techniques are more explicitly combined with participatory science processes, like the GFSC, the scale of information and knowledge that could be gained would be instrumental. In addition to data collection and knowledge transfer, other benefits would include increased ocean literacy, acceptance of the need for management and policy change, and the capacity of communities to work together to understand and tackle conservation challenges (Hind-Ozan *et al*. 2017). However, some caveats are needed here as well. Scalable innovative solutions are needed to more efficiently gather, process, analyze, and distribute observational data from across users, but users also need to be empowered to control their data. Even in the GFSC, individuals did not want to share the exact location of sites for fear of their sharks being fished out. Therefore, data ethics and the ability of users to maintain their data rights (e.g., self-determination, self-declaration) will also be important.

For the GFSC, the scientists at eOceans manually processed and analyzed the data. However, eOceans used the lessons learned throughout this study and other similar projects (e.g., eShark, eManta, Global Shark Sanctuary Assessment) to build a mobile app and cloud-based platform that digests, processes, maps, and analyzes data similar to the GFSC in real-time. It works for tracking of all marine species (>200,000 species listed), activities, and issues (e.g., pollution, injuries, entanglements, diseases) in all areas of the ocean (based on a user supplied area boundary file) logged by any activity type. To facilitate data collection and local knowledge transfer, eOceans added an in-app field guide that helps data contributors ensure they are in agreement on the species being recorded, and the co-interpretation channel was designed for contributors to view the live results of the project and provide comments regarding their interpretations. The eOceans platform enables any scientist, researcher, or community group to repeat the GFSC assessments for any other species of interest in any part of the ocean. Instead of data contributors having to write their observations down in community logbooks, as with the GFSC, they are empowered to log observations in the mobile app where they own and have access to their data and decide which projects they choose to share their data with, thus empowering data democracy and self-determination. They can choose to share with a few or hundreds of projects by simply joining the project. The analyses displayed in the current study for the GFSC are being integrated into the eOceans platform for real-time tracking of effort and observations, so that the GFSC and others can be resumed to continuously track sharks and other species in real-time.

## Conclusions

By aggregating divers’ observations from across Fiji, the GFSC created an extensive longitudinal dataset on hundreds of dive sites and enabled the first account of where eleven shark species spend at least part of their time and added new insights on their distribution and changes to relative abundance. Although there are limitations associated with this type of data, the extent of sampling and the reliability of results demonstrate the value of collaborative co-generated programs for filling data and perspective gaps on marine fauna, and the power of a community to come together to assess the issues that matter to them.

These results may be used to prompt new scientific questions, to evaluate or resolve management and policy strategies, or as a contemporary baseline that future shark populations or human use patterns may be compared against. As well, since participatory science and co-generated science programs of this magnitude considerably increase the number of opportunities to exchange knowledge, experience, and ideas between industry and science, while also providing frequent touch points with the broader community (e.g., tourists) to promote education and outreach opportunities on issues affecting the oceans, they have many added benefits. These social outcomes may help meet various sustainable development and biodiversity conservation goals, and deserve further inclusion in scientific and management discussions.

## Data Availability

The data are held by the GFSC team in Fiji and stored in the eOceans data collection and management platform, and are available from either on reasonable request.

