## Supplemental Figures for "Community-driven shark monitoring for informed decision making: A case study from Fiji"

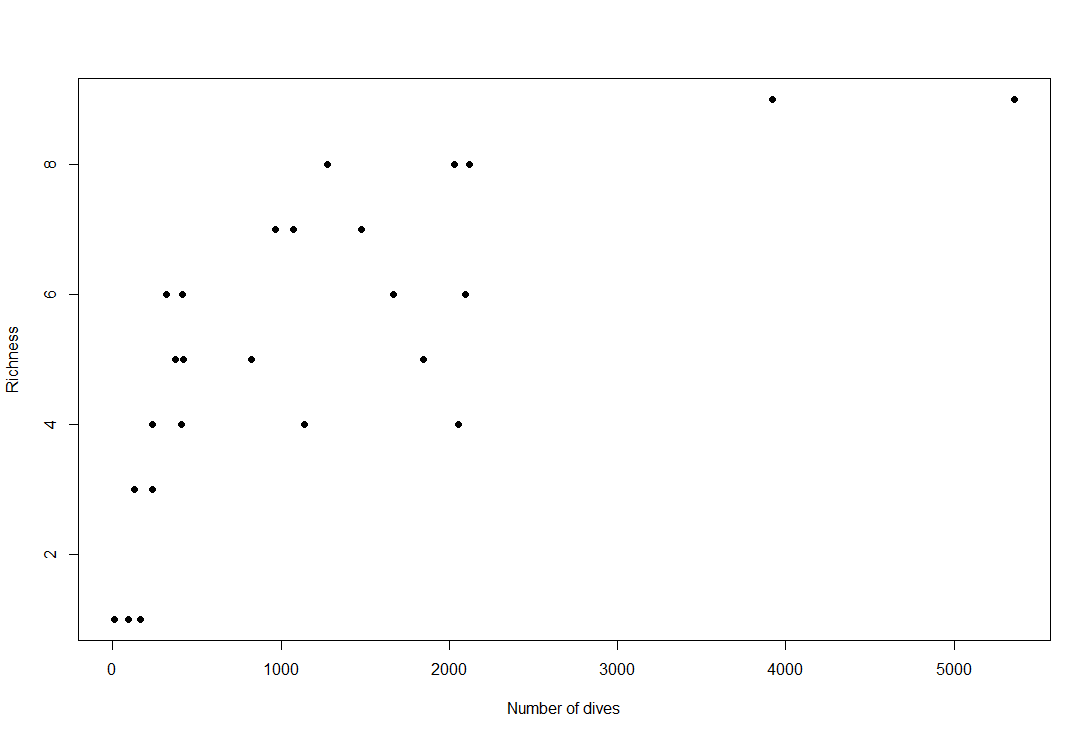

**Figure S1. Species accumulation curve by effort.** Total shark species richness observed in an area compared to total effort in number of dives that occurred in that area.

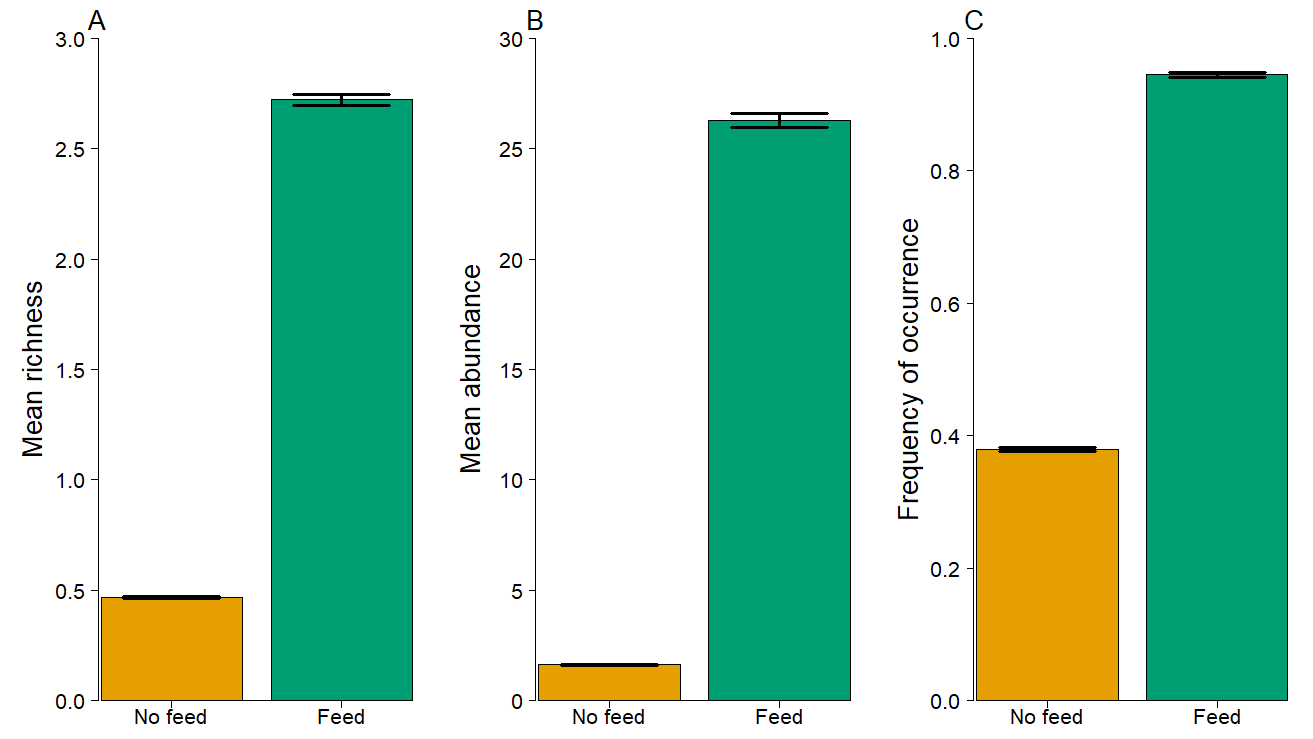

**Figure S2. Sharks at feeding and non-feeding sites.** (A) mean species richness, (B) mean abundance, and (C) frequency of occurrence at non-feeding (orange) versus feeding (green) dives. Bars are ± SE.

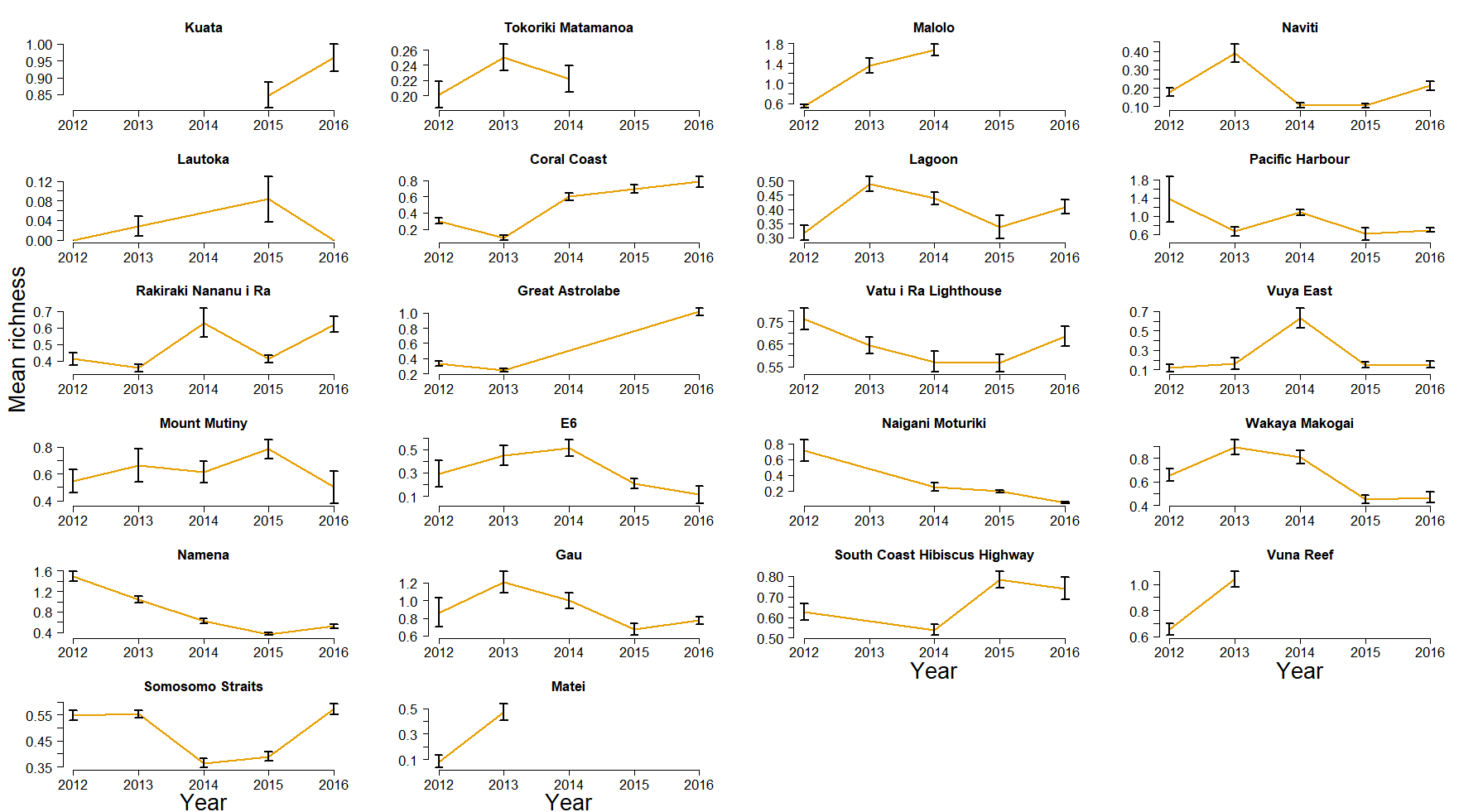

**Figure S3. Mean richness (± SE) of sharks in each area at non-feeding sites across each year, excluding areas surveyed in only one year.** Areas are ordered left to right and top to bottom from most western to most eastern.

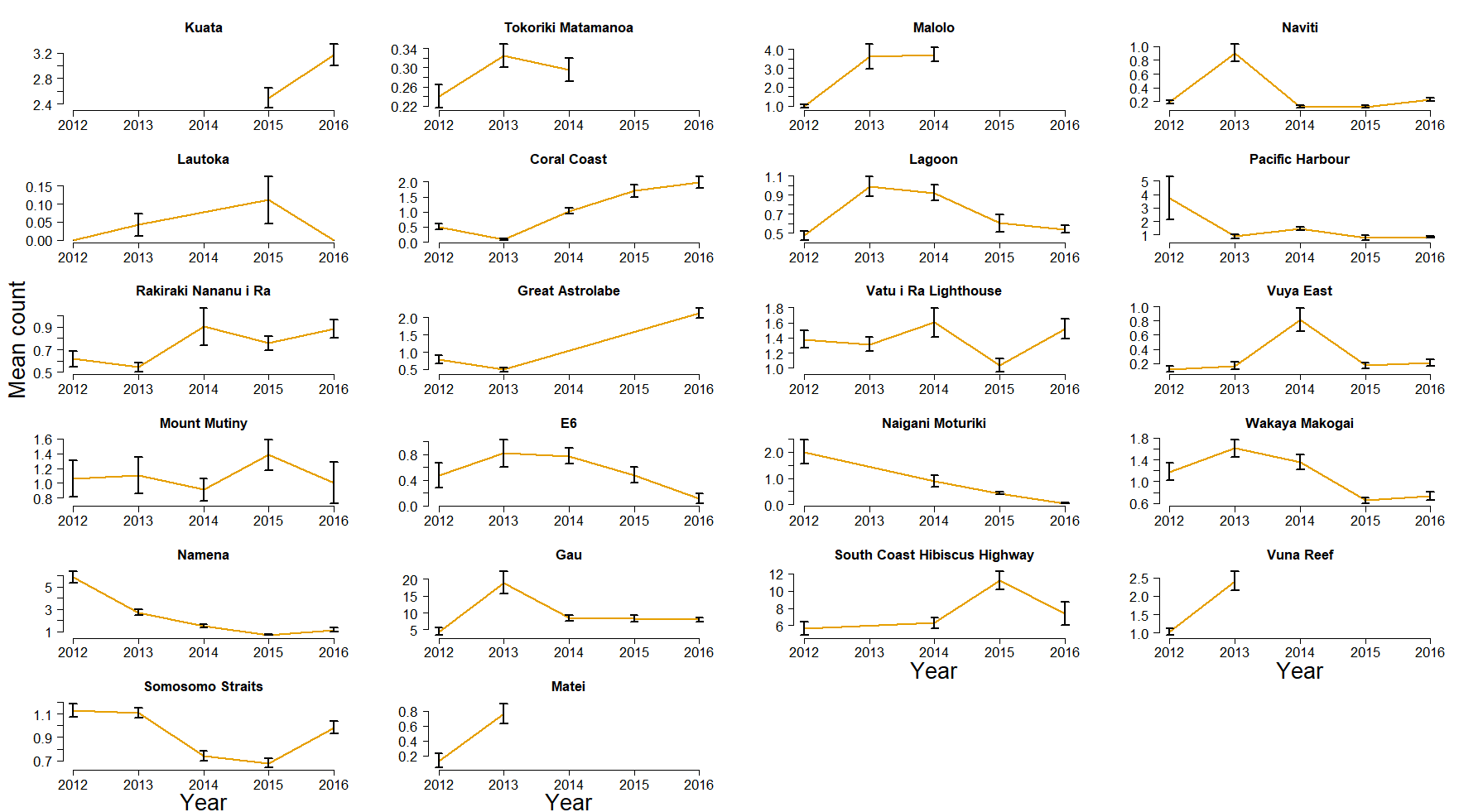

**Figure S4. Mean abundance (± SE) of sharks in each area at non-feeding sites across each year excluding areas surveyed only in one year.** Areas are ordered left to right and top to bottom from most western to most eastern.

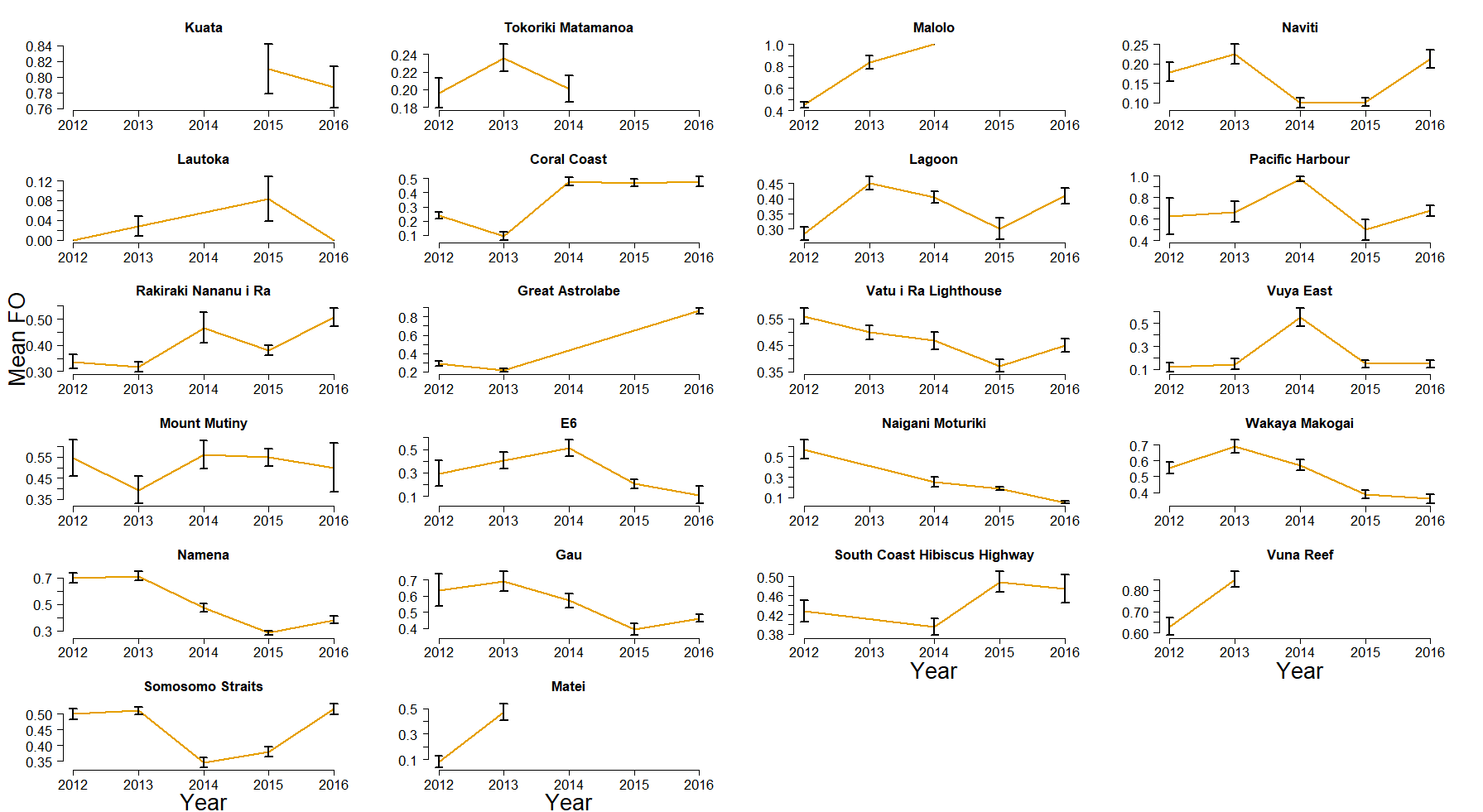

**Figure S5. Frequency of occurrence (± SE) of sharks in each area at non-feeding sites across each year excluding areas surveyed only in one year.** Areas are ordered left to right and top to bottom from most southern to most northern.

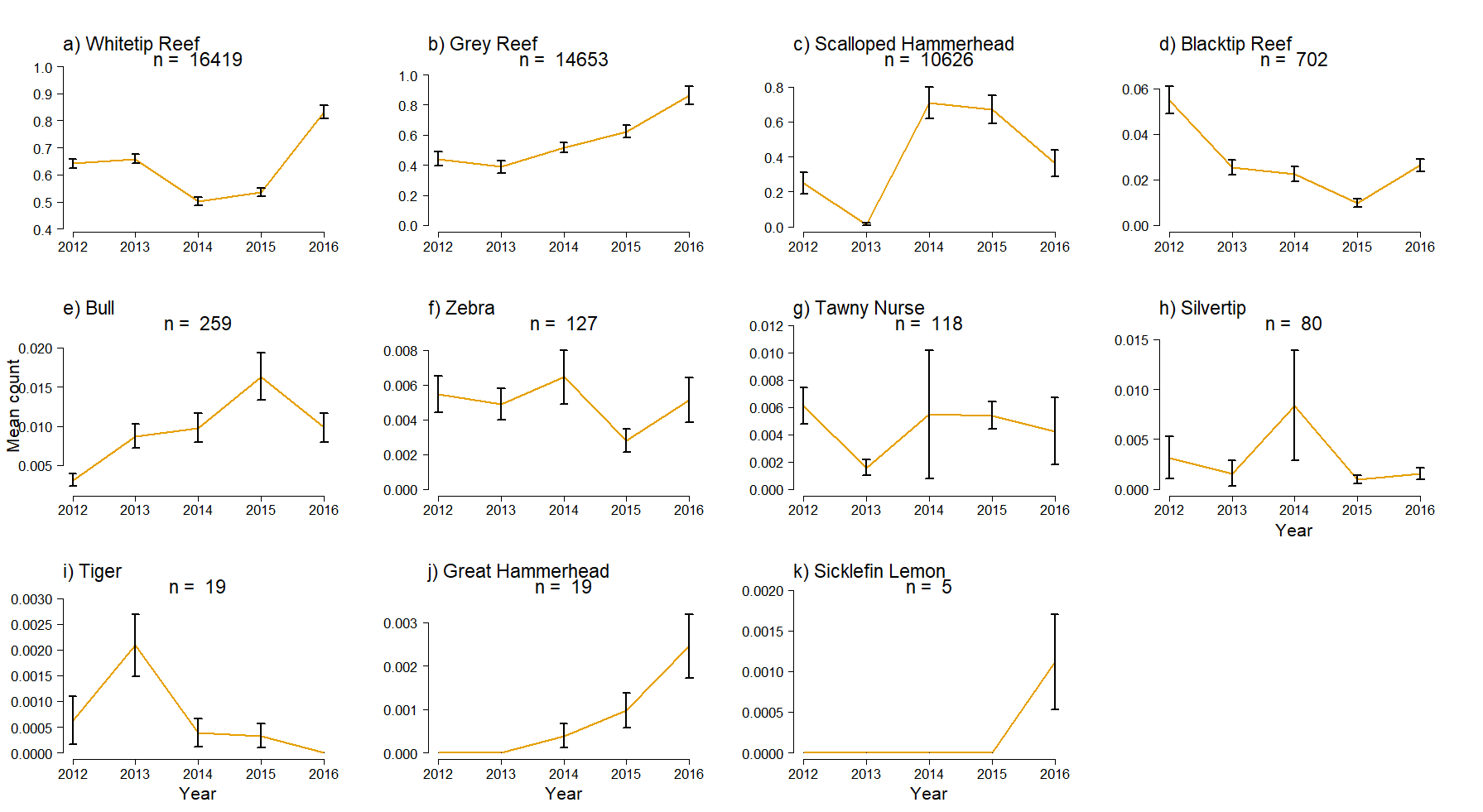

**Figure S6. Variability in abundance by species through time by species at non-feeding sites.** The mean count (± SE) at non-feeding sites across areas visited in more than one year (n=22), ordered by total count (numbers on top) from left to right and top to bottom.

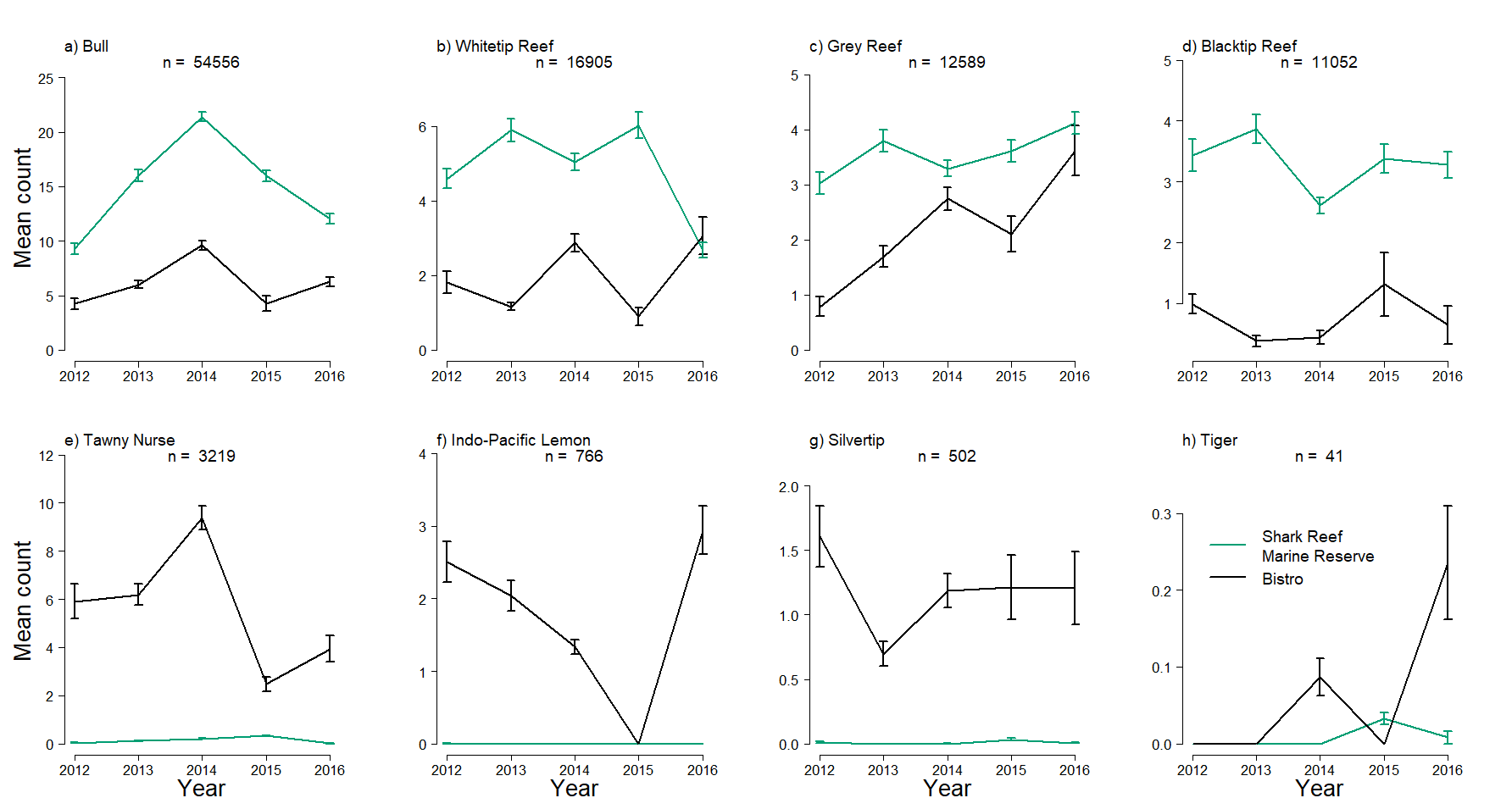

**Figure S7. Annual variability of sharks by species at feeding sites**. The mean count (± SE) of each species at the two most visited feeding sites in Pacific Harbour across each year, ordered by total abundance from left to right and top to bottom.
